# Chronic social stress blunts core body temperature and molecular rhythms of *Rbm3* and *Cirbp* in mouse lateral habenula

**DOI:** 10.1101/2023.01.02.522528

**Authors:** Salma Haniffa, Priyam Narain, Michelle Ann Hughes, Aleksa Petković, Marko Šušić, Vongai Mlambo, Dipesh Chaudhury

**Author notes:** These authors contributed equally to this work (Salma Haniffa, Priyam Narain).

## Abstract

Chronic social stress in mice causes behavioral and physiological changes that result in perturbed rhythms of body temperature, activity and sleep-wake cycle. To further understand the link between mood disorders and temperature rhythmicity in mice that are resilient or susceptible to stress, we measured core body temperature (Tcore) before and after exposure to chronic social defeat stress (CSDS). We found that Tcore amplitudes of stress-resilient and susceptible mice are dampened during exposure to CSDS. However, following CSDS, resilient mice recovered temperature amplitude faster than susceptible mice. Furthermore, the interdaily stability (IS) of temperature rhythms was fragmented in stress-exposed mice during CSDS, which recovered to control levels following stress. There were minimal changes in locomotor activity after stress exposure which correlates with regular rhythmic expression of *Prok2* - an output signal of the suprachiasmatic nucleus. We also determined that expression of thermosensitive genes *Rbm3* and *Cirbp* in the lateral habenula (LHb) were blunted 1-day after CSDS. Rhythmic expression of these genes recovered 10 days later. Overall, we show that CSDS blunts Tcore and thermosensitive gene rhythms. Tcore rhythm recovery is faster in stress-resilient mice, but *Rbm3* and *Cirbp* recovery is uniform across the phenotypes.

## Introduction

Daily cycles of light and temperature are the two primary environmental timing cues used by living systems to entrain their endogenous circadian rhythms to the astronomical day. In mammals, the suprachiasmatic nucleus (SCN), the principal biological clock, drives circadian rhythms of core body temperature (Tcore), which is maintained within a narrow range to allow for optimal physiological functioning [1,2]. These rhythms can be entrained by external cues, predominantly light, but in the absence of these cues they exhibit endogenous free running characteristics. In homeothermic organisms Tcore is measured by internal and peripheral thermoreceptors [3]. The preoptic area of the hypothalamus is the primary integrative site for thermoregulation [4]. The narrow range of Tcore rhythms is maintained by a complex feedback system consisting of heat-loss mechanisms that are activated when Tcore temperature rises, as well as heat-gain mechanisms which are activated when it falls [5–7]. It is hypothesized that daily Tcore rhythms are generated by SCN neurons projecting into hypothalamic regulatory centers and by modulation of thermogenic brown adipose tissue [8]. Tcore is an intrinsic zeitgeber (ZT) since changing the amplitude and frequency of temperature rhythms affects the rhythmic expression of key clock genes: Basic Helix-Loop-Helix ARNT Like 1 (*Bmal1*) and Period 1 (*Per1*) [9]. These changes are likely driven by changes in expression of thermo-sensitive genes, such as RNA-binding Motif Protein 3 (*Rbm3*), Cold-inducible RNA binding protein (*Cirbp*) and Heat Shock Factor 1 (*Hsf1*), which are chaperones that modulate expression of a variety of target genes including the clock genes [7,10–13].

Mood disorders and aberrations in Tcore rhythms are closely related. Patients often exhibit a shift or blunting of diurnal cortisol and Tcore rhythmic activity during depressive episodes [14,15]. The association between circadian rhythms and depression is commonly attributed to molecular and pathophysiological changes in brain circuits that coregulate emotions, locomotor activity, sleep and body temperature [16]. Although clinical studies indicate a relationship between desynchronization of Tcore rhythms and symptom severity, it is unclear whether the relationship is correlational or causal [15]. While the effects of stress on activity, sleep-wake cycle, and molecular rhythms have been investigated, mechanisms linking stress and Tcore rhythms are understudied.

Stress-induced thermogenesis of brown adipose tissue that results in emotional hyperthermia is thought to be driven by the lateral habenula (LHb) and ventral tegmental area (VTA) [17,18] – two key brain regions that exhibit elevated pathophysiological firing in stress-susceptible mice [19–22]. Pathophysiological changes in these regions may be responsible for stress-induced changes in temperature rhythms. Furthermore, onset of a variety of mood disorders has been associated with changes in circadian rhythms of molecular processes in the SCN and LHb [20,23]. At the transcriptome level, *Rbm3* and *Cirbp* are expressed in response to lower temperatures while *Hsf1* is expressed in response to heat stress [24]. Patients with mood disorders exhibit a higher expression of *Rbm3* compared to healthy controls [25]. Deletion of *Rbm3* leads to a significant reduction in expression amplitudes of clock genes *Bmal1*, *Clock*, *Per1* and *Cry1* Cryptochrome 1(*Cry1*) [10]. In the cell nucleus, *Cirbp* binds to the 3’-UTR of *Clock* mRNA and promotes its translocation to the cytoplasm, thereby enabling its translation [11–13]. *Hsf1* interacts with clock genes, such as *Bmal1,* in order to maintain synchronization of the cellular clock and clock-controlled adaptive responses to cellular stress [26]. Absence of *Hsf1* activity can lead to impaired hippocampal development and severe behavioral disruptions, such as depression and elevated aggression [27].

To understand the association between Tcore and mood disorders we used the chronic social defeat stress (CSDS) paradigm to investigate the effects of chronic stress on Tcore and expression of *Cirbp*, *Rbm3* and *Hsf1* (Fig. 1). Prior to stress all mice exhibited similar daily Tcore rhythms. During CSDS both stress-resilient and susceptible mice exhibited blunted Tcore rhythms. During the first 5 days of post-CSDS recovery period, Tcore amplitudes were significantly blunted in susceptible, but not resilient, mice compared to controls. By days 6-10 of post-CSDS recovery, the average Tcore amplitude of susceptible mice returned to control levels (Fig. 2). Furthermore, stress-exposed mice exhibited fragmented temperature rhythms during CSDS, which was consolidated in the recovery phase. Control mice exhibited daily rhythmic expression of *Rbm3* and *Cirbp* in the LHb where transcript levels were higher in the day than the night – in concordance with decreased core body temperature in the day compared to the night (Fig. 3). In contrast, rhythmic expression of *Rbm3* and *Cirbp* expression was blunted 1-day post-CSDS in both resilient and susceptible mice which returned to control levels after 10 days of recovery. *Hsf1* did not exhibit diurnal rhythmic expression in the LHb of control mice and was not impacted by stress exposure (Supplemental Fig.5). *Prok2* expression in the SCN was unaffected by stress exposure as all mice exhibited high relative expression in the day and low at night which likely correlates with normal locomotor rhythms in mice exposed to stress (Supplemental Fig. 3, Supplemental Fig. 4). *BMAL1* expression likewise did not differ between the phenotypes (Supplemental Fig.4).

**Figure 1.**
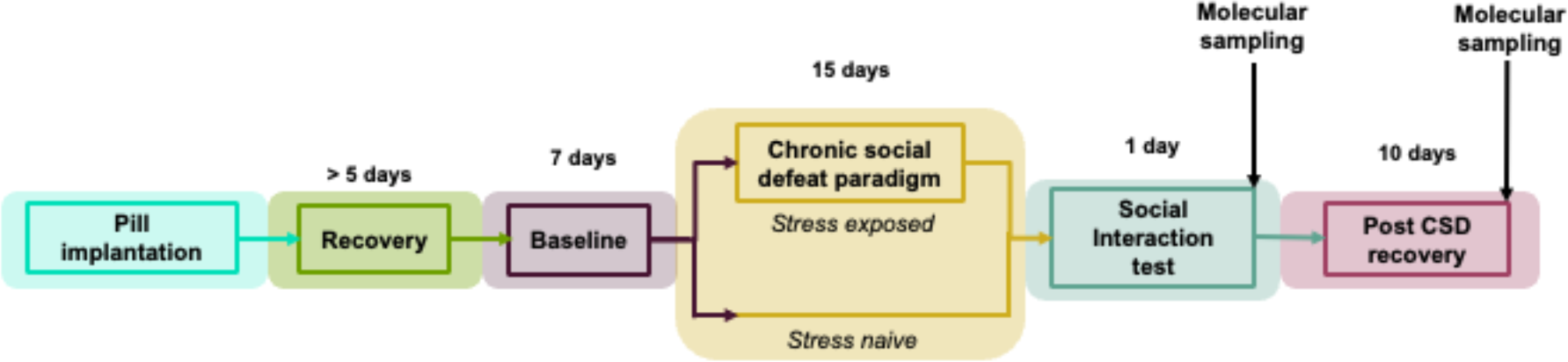
Timeline for investigating the association between stress, diurnal temperature rhythms and changes in expression of thermosensitive genes.

**Figure 2.**
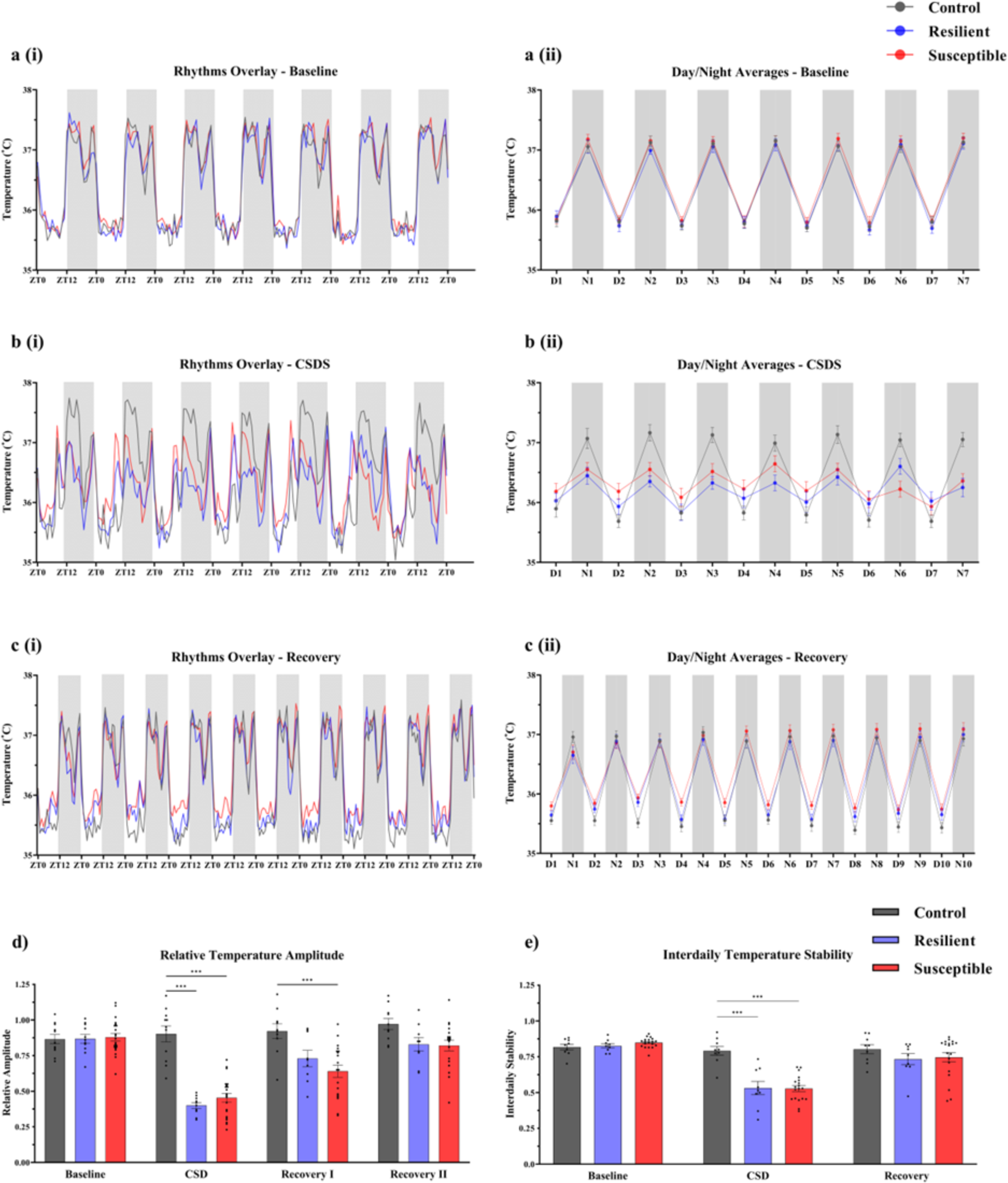
Average core body temperature rhythms during baseline, CSDS and recovery. Temperature values include recordings from 7 days of baseline, last 7 days of CSDS and 10 days of recovery. **a(i)** Control, resilient and susceptible mice show comparable Tcore rhythms prior to CSDS in 12H:12H light/dark conditions. **b(i)** CSDS blunts the temperature amplitude of resilient and susceptible mice compared to controls (N=11). **c(i)** The Tcore of susceptible mice remains blunted longer than resilient mice during the recovery period. **a-c (ii)** Average day and night values (averaging 12h day and night separately for each phenotype) of Tcore during baseline, CSDS and recovery. **Baseline:** No significant difference between phenotypes. **CSDS:** Susceptible and resilient mice exhibit significantly lower Tcore most nights compared to controls. Two-way ANOVA revealed significant main effect of phase (F_13,462_ = 21.90, *p* < .001), phenotype (F_2,462_ = 11.36, *p* < .001), and interaction between the two factors (F_26,462_ = 4.738, *p* < .001). In most nights, resilient and susceptible mice exhibited significantly lower average Tcore compared to controls (Supplemental Table 1). **Recovery:** Daytime Tcore of susceptible mice was significantly higher relative to controls for most days following CSDS. Two-way ANOVA shows significant effects of phase (F_19,660_ = 148.7, *p* < .001) and phenotype (F_2,660_ = 20.08, *p* < .001). Daytime Tcore averages of susceptible mice during recovery were higher on most days relative to controls (Supplemental Table 2). **(d) Baseline**: There was no difference in baseline Tcore amplitude between the phenotypes. **CSDS**: Tcore amplitude was significantly blunted in susceptible and resilient mice during CSDS relative to controls (Supplemental Table 2). **Recovery**: Susceptible and resilient mice exhibited significantly blunted amplitude relative to the controls during the first 5 days of recovery. Temperature amplitude recovers to control levels in days 6-10. **(e)** Interdaily stability of the temperature rhythms during baseline, CSDS and recovery were calculated for each phenotype. Two-way ANOVA shows significant effects of phase (F_2,64_ = 62.3, *p* < .001), phenotype (F_2,34_ = 6.24, *p* < .01) and interaction (F_4, 68_ = 11.9, *p* < .001). **Baseline**: There was no difference in Tcore rhythm stability between the phenotypes. **CSDS**: Susceptible (Tukey’s HSD, *p* < .001, 95% C.I. = [0.17, 0.36]) and resilient mice (*p* < .001, 95% C.I. = [0.11, 0.40]) exhibit significantly lower Tcore rhythm stability relative to controls. **Recovery**: There was no difference in Tcore rhythm stability between the phenotypes. (a-e) Sample sizes: control (*N* = 9-11), resilient (*N* = 9-11), and susceptible mice (*N* = 19-23). Error bars represent mean ± SEM. *** indicate *p* < .001.

**Figure 3.**
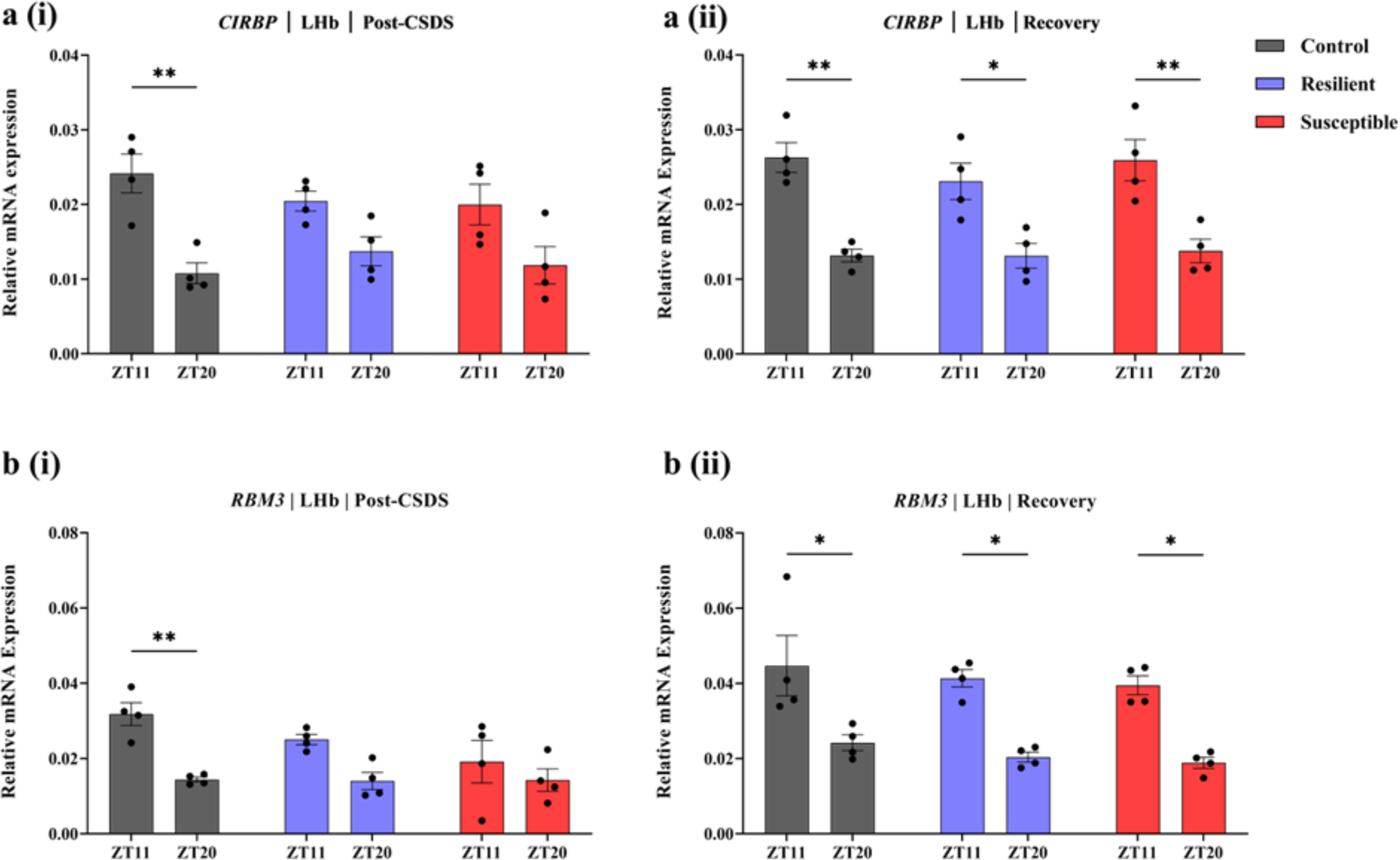
Diurnal rhythmic expression of *Cirbp* and *RBM3* in the LHb is blunted immediately after CSDS. a(i) & b(i) Control, but not resilient and susceptible mice, display significantly larger mRNA expression of *Cirbp* (Tukey’s HSD, *p* < .01, 95% C.I. = [0.00, 0.02]) and *RBM3* (*p* < .01, 95% C.I. = [0.00, 0.03]) in the LHb at ZT11 compared to ZT20 post-CSD. **a(ii) & b(ii)** The molecular rhythms of *Cirbp* and *RBM3* in resilient [*Cirbp: p* = .02, 95% C.I. = [0.00, 0.02]; *RBM3: p* = .01, 95% C.I. = [0.00, 0.04]] and susceptible [*Cirbp: p* < .01, 95% C.I. = [0.00, 0.03]; *RBM3*: *p* = .01, 95% C.I. = [0.00, 0.04]] mice return 10 days after recovery and are at similar levels comparable to control [*Cirbp: p* < .01, 95% C.I. = [0.00, 0.02]; *RBM3: p* = .01, 95% C.I. = [0.00, 0.04]]. Each experimental group consisted of 4 biological replicates split into 3 technical replicates (*N* = 4). Error bars represent mean ± SEM.* and ** indicate *p* < .05 and *p* < .01, respectively.

## Methods

### Ethics statement and animals

Animal protocols have been approved by the National Institute of Health Guide for Care and Use of Laboratory Animals (IACUC Protocol: 150005A2, 19- 0004A1), as well as the NYUAD Animal Care and Use Committee. C57BL6/J mice of 5-7 weeks old were purchased from Jackson Laboratory (Bar Harbor, Maine, USA). Single-housed male CD1 retired breeder mice from Charles River laboratory (Wilmington, Maine, USA) were used as resident aggressors for the CSDS paradigm. All experimental mice were maintained within standard housing conditions at a temperature of 21±2°C and humidity of 50±10% with *ad libitum* access to food and water, and enrichment in the form of wood shavings. Mice were kept on a 12:12-hour light/dark cycle (light onset at 7am (ZT0), light offset at 7pm (ZT12)). Temperature, locomotor and gene expression experiments were done on the same mouse cohorts.

### Core body temperature recordings

Mice were anesthetized with a ketamine (100 mg/kg) and xylaxine (10 mg/kg) mixture. A wireless electronic telemetry pill (Anipill; BodyCap, Paris, France) was surgically implanted into the peritoneal cavity of all experimental mice. This surgical implantation was carried out by adopting and modifying previously published protocol [28]. The Anipill system was configured to collect temperature measurements every 15 minutes. Temperature recordings were set to begin immediately after pill activation and continued to collect and transmit data for the entire duration of the experiment. Open-source online software Biodare2 was utilized to extract circadian parameters such as amplitude, phase, and period of temperature rhythms [29]. Fast Fourier transform non-linear least squares (FFT NLLS) was employed to analyze average hourly data. Interdaily stability analysis was conducted in R (version 4.1.1) following previously published equations [30,31].

### Locomotor activity recording

Locomotor activity during baseline and recovery periods was continuously measured using Actimetrics ClockLab wireless passive infrared (PIR) sensor system (Actimetrics, Chicago, Illinois, USA). The infrared sensor was placed on top of each small hamster cage, which were used to single-house the mice. The data was collected using ClockLab Data Collection Software and analyzed with ClockLab Analysis Software (version 6.1.10).

### Chronic social defeat stress and behavioral analysis

For a period of 15 days, experimental C57BL/J6 mice were exposed daily to an aggressive CD1 resident mouse for 10 minutes of physical interaction. After each defeat session, C57BL/J6 mice were separated from the CD1 mice within the same cage using a perforated plexiglass divider that allowed for olfactory and visual sensory, but not physical, contact. Daily defeat sessions were consistently carried out between ZT8 and ZT10. C57BL/J6 mice were exposed to a novel aggressor each day to avoid familiarization. Control mice were kept in pairs in an equivalent cage set-up, with a plexiglas divider separating the two control mice at all times. These mice were kept in the same room as experimental mice in order to control for variations in ambient temperature.

To assess social aversion as a measure of depressive-like behavior, we utilized the social interaction (SI) test. For each cohort, the SI test was conducted the day following the end of CSDS between ZT6 and ZT10. The test was conducted in a white square plexiglass arena (dimensions 42 X 42 X 42.5cm) under red light. The SI test for each mouse consisted of two 2.5minute sessions: one without a CD1 social target and another with an unfamiliar nonaggressive CD1 social target. TopScan was used to track the time each experimental mouse spent in the predefined zones within the arena. Mice were phenotyped based on the social interaction ratio (SI ratio) scores, which was calculated as described previously [32]. Experimental mice with scores < 100 were classified as ‘stress-susceptible’ and those with scores ≥ 100 were classified as ‘stress-resilient’.

### Brain sampling for molecular analysis

Sampling was started 24 hours following SI and completed within three days. Mice were anesthetized using 0.25ml isoflurane (Vedco, St. Joseph, Missouri, USA) and then sacrificed by cervical dislocation followed by decapitation. Extractions were carried out on a cold sterile surface and the extracted brains were directly immersed into cold isopentane (Sigma Aldrich M3263) to be flash-frozen. Brains were sliced on the Leica CM1950 cryostat (Leica Biosystems, Nussloch, Germany), using Mouse Brain Atlas [33] as a reference for identification of the SCN and the LHb. Brain slices were visualized under Leica S Apo Stereozoom 1.0x-8.0x stereoscope (Leica Microsystems, Heerburg, Switzerland) and tissue punches of the regions were collected into 1.5ml DNA LoBind tubes (Eppendorf, Hamburg, Germany). The punches were then snap-frozen using dry ice before being stored at -80 C° until RNA isolation.

### RNA isolation and quantitative PCR

RNA was isolated using Qiagen RNeasy Micro Kit, as per manufacturer’s instructions, and quantified using Nanodrop 8000 (Thermo Fisher Scientific, Waltham, Massachusetts, USA). The isolated RNA was converted to cDNA using the Maxima H Minus First Strand cDNA Synthesis Kit (Thermo Fisher Scientific), as per manufacturer’s protocol. The template cDNA was amplified using Qiagen Primers for the transcripts: Hsf1 (NM_008296), Rbm3 (NM_016809), Cirbp (NM_007705) and Prok2 (NM_015768) to quantify levels of mRNA in the SCN and LHb. GAPDH (NM_008804) was used as the reference housekeeping gene in both SCN and LHb, as reported previously [34–39]. Non-template controls were run for each primer pair. qPCR was carried out using Applied Biosystems QuantStudio5 Real-Time PCR System (Thermo Fisher Scientific). All experimental groups for qPCR experiments consisted of 4 biological replicates split into 3 technical replicates to ensure accuracy of measurement. The average of the threshold cycle (Ct) was taken across the triplicates and normalized to the GAPDH using the 2^(ΔCt)^ method. Fold changes were calculated relative to controls using the 2^(ΔΔCt)^ method.

### Statistical analysis

Prism 9 (Graphpad Software, La Jolla, California, USA) was used for data plotting and statistical analyses. Statistical significance was determined using one-way and two-way ANOVAs, two-tailed independent *t*-tests and Tukey’s HSD *post hoc* test. All values are expressed as the mean ± standard error of the mean (SEM). The sample size (*n*) is noted in the specific figure legends for each experiment.

## Results and Discussion

We found that daily rhythms of Tcore are affected by chronic social stress. Prior to stress, all mice exhibited lowest Tcore in the daytime, when mice are least active, and highest at nighttime, where they are most active (Fig. 2a(i,ii)) (Supplemental Table 1). However, during CSDS both resilient and susceptible mice generally exhibited higher Tcore in the daytime and lower Tcore in the nighttime, compared to controls (Fig. 2b(i,ii)). Daytime Tcore of resilient mice returned to control levels immediately after CSDS, while susceptible mice continued to exhibit elevated Tcore for a longer period during the recovery phase (Fig. 2c(i,ii)). During CSDS, both resilient and susceptible mice exhibited significantly lower Tcore amplitudes compared to controls (Fig. 2d) (Supplemental Table 2). However, during the first 5 days of recovery period, Tcore amplitudes were significantly blunted in susceptible, but not resilient mice. In contrast there were no significant differences in Tcore amplitude between control and susceptible from day 6 to day 10 of the recovery period. Furthermore, the interdaily stability (IS) of temperature rhythms during stress was fragmented in all groups which recovered to control levels post stress (Fig. 2e). Our results align with clinical findings that patients with depression exhibit blunted diurnal rhythms where nighttime temperature is abnormally high (when people are usually least active), while daytime temperature is largely unaffected [15]. Preliminary studies in humans suggest that circadian temperature profile predicts stress vulnerability [40–42]. However, since there were no differences at baseline in our study, we failed to model these results using CSDS. Previous studies have shown aberrations in physiological processes in susceptible mice [19,43–45]. It is likely that homeostatic factors that maintain Tcore in resilient mice buffer against stress-induced processes that drive elevated daytime Tcore and greater blunting of Tcore amplitudes in susceptible mice. Our study correlates with previous observations that more severe stress induces significantly greater blunting of temperature rhythms [46]. We had previously shown that susceptible and resilient mice display reduced overall sleep homeostasis [47]. Moreover, susceptible mice exhibited deficient NREM recovery sleep responses and increased NREM sleep fragmentation following CSDS [48,49]. Although affected by locomotion and sleep patterns, Tcore is under tight circadian control which exerts the most powerful influence on its regulation [50]. Since sleep and Tcore are closely related it may be that chronic stress causes pathophysiological changes in circuitry that coregulates sleep architecture and body temperature homeostasis [51,52].

Given the regulatory role of the SCN in governing temperature rhythms, we speculated that additional rhythmic processes influenced by the SCN might also be perturbed by chronic stress. Thus, we recorded locomotor rhythms before and after exposure to CSDS and found these rhythms to be unaffected by chronic stress (Supplemental Fig. 3). The three phenotypes exhibited similar bouts of activity, light and dark phase activity profiles, as well as interdaily and intradaily stability of locomotor rhythms ((Supplemental Fig. 3)). We did not observe any clear correlation pattern between Tcore and locomotor activity in any of the phenotypes or recording phases (Supplemental Fig.8). Since we did not observe any effect of stress on locomotor rhythms, we wondered whether this lack of effect was associated with expression patterns of genes that regulate diurnal cycles of locomotion. Prokineticin 2 (*Prok2*) is one such clock-controlled gene whose product is an important neuropeptide output signal of the SCN that regulates locomotor rhythms [53]. Relatively stable expression of *Prok2* in the SCN of stress-exposed mice, along with rhythmic diurnal locomotor activity, suggests that the SCN is buffered against environmental stressors, enabling normal light entrainment (Supplemental Fig. 4 a(i,ii)). In line with this hypothesis, we observed similar *BMAL1* expression patterns in the SCN across the three phenotypes (Supplemental Fig. 4 b(i,ii)). Relatedly, clock gene expression following chronic mild stress (CMS) was shown to be stable in the SCN but not in the other regions involved in mood regulation [54]. Previous light pulse experiments on stress-exposed and control mice revealed phase delay of the body temperature rhythm without any changes in the circadian pacemaker functions [55]. Our findings on locomotor activity and *Prok2* expression match previous studies and indicate that pacemaker function remains unaffected by different forms of chronic stress (Supplemental Fig. 3, Supplemental Fig.4) [54,55].

Next, we investigated whether expression of the thermosensitive genes *Rbm3*, *Cirbp* and *Hsf1* correlates with the observed changes in temperature profiles immediately after CSDS and 10 days of recovery. *Rbm3* and *Cirbp* are both expressed in response to lower temperatures, with their baseline expression oscillating diurnally and being linked to circadian function [7,56,57]. The expression of *Rbm3* and *Cirbp* in the LHb of control mice was significantly higher in the day (ZT11), when Tcore is lower, compared to the night (ZT20), when Tcore is higher (Fig. 3ai,bi). In contrast, immediately after CSDS, there was no significant difference in daytime and nighttime expression of *Rbm3* and *Cirbp* in the LHb of susceptible and resilient mice. However, after 10 days of recovery, the diurnal differences in expression of *Rbm3* and *Cirbp* in the LHb of both susceptible and resilient mice were similar to control mice (Fig. 3aii,bii), suggesting recovery of daily rhythmic expression following the recovery of temperature rhythms. *Hsf1* did not show robust rhythms in the LHb (Supplemental Figure 5). Studies have shown that exposure to decreased temperature (cold shock) does not generate a significant expression of *Hsf1*, consistent with the findings from our study [58]. Additionally, the lack of significant changes within the LHb suggest possible region-specific differences in rhythmic expression following exposure to chronic stress (Supplemental Fig. 5).

These results are supported by previous findings that mild hypothermia results in peak expression of *Cirbp* and *Rbm3* which decreases significantly following deep hypothermia [12]. In turn, hyperthermia causes substantial decreases in expression of both of these genes in cultured mammalian cells [12]. CMS affects rhythmic expression of clock genes in various brain regions, including the LHb [54]. We suggest that chronic stress disrupts Tcore resulting in perturbations of thermosensitive gene expression which then impacts clock gene expression in the long-term. This eventually results in pathophysiological changes in the LHb [21]. Since we did not see any stress induced changes in clock-controlled gene expression we do not think stress directly disrupts thermosensitive gene expression. To the best of our knowledge, the selected thermosensitive genes do not affect temperature regulation directly. Additionally, the identical pattern of blunting observed in core body temperature and expression of these genes in the LHb at the same time points further suggests that it is core body temperature that causes this gene expression blunting rather than an independent CSDS-induced mechanism. Further experiments involving higher temporal resolution and examination of additional clock genes are necessary, to support our conclusion. Further experiments involving higher temporal resolution and examination of additional clock genes are necessary, to support our conclusion. Our observations on the effect of stress on temperature and molecular rhythms are in close agreement with previous findings where repeated 5-day social defeat stress resulted in decreased amplitude of heart rate and body temperature persisting for at least 10-days after the final confrontation [59]. *Rbm3* regulates local synaptic translation and neuronal activity, and knockdown of *Rbm3* at specific times of the day increases firing of cultured hippocampal cells [57]. Increased expression of *Cirbp* when mice are asleep, possibly because of lower Tcore associated with lower motor activity, attenuates expression of wake-inducing/sustaining genes, such as *Homer.* In contrast, sleep deprivation leads to increased expression of wake-inducing genes because of prolonged attenuation of *Cirbp* expression, possibly because of elevated Tcore as a result of greater activity [60]. We speculate that such evidence points to the mechanism by which stress induces fragmented sleep resulting in decreased *Cirb* expression and, consequently, greater expression of wake-promoting genes that cause further sleep fragmentation. Future studies are warranted to investigate this putative feedforward mechanism and its role in interlinking mood disorders, fragmented sleep and Tcore changes.

## Conclusion

Taken together, this study provides additional evidence for stress-induced changes in temperature regulation and differential expression of thermo-sensitive genes. Importantly, we identified physiological differences in daily temperature profiles between susceptible and resilient mice. As such, this study represents a crucial step towards a more systematic understanding of the temperature imbalances experienced by patients diagnosed with mood disorders. The preliminary findings of this study highlight the interplay between temperature dysregulation and thermosensitive gene expression in the LHb. One limitation of this study is the lack of multiple sampling timepoints across the day/night cycle that would better assess rhythmic gene expression with higher resolution. Future studies are warranted to include more timepoints and investigate rhythmicity of expression of other clock genes in response to stress. Some genes may appear relatively unaffected at the mRNA levels but their protein levels may be altered. Future experiments to investigate the changes in protein expression would enable an additional level of insight into the effects of stress on core body temperature regulation. Moreover, investigation to determine if there are instances where changes in Tcore increase stress vulnerability are also warranted. Overall, such studies would help clarify the cause and consequence in the interplay between core temperature and clock-modulating thermosensitive genes.

## Abbreviations

(BAT): Brown adipose tissue
(Bmal1): Basic Helix-Loop-Helix ARNT Like 1
(Cirbp): Cold inducible RNA binding protein
(CMS): Chronic mild stress
(CSDS): Chronic social defeat stress
(FFT NLLS): Fast Fourier Transform Non-Linear Least Squares
(Hsf1): Heat Shock Factor 1
(LHb): Lateral habenula
(Per1): Period 1
(PIR): Passive infrared
(POA): Preoptic area
(Rbm3): RNA- binding Motif Protein 3
(Tcore): Core body temperature
(SCN): Suprachiasmatic nucleus
(VTA): Ventral tegmental area

## Author contributions

**Conceptualization:** Vongai Mlambo, Salma Haniffa, Michelle Ann Hughes, Priyam Narain, Dipesh Chaudhury

**Data curation:** Salma Hanniffa, Michelle Ann Hughes

**Formal analysis:** Salma Haniffa, Michelle Ann Hughes, Marko Šušić, Aleksa Petković, Priyam Narain, Dipesh Chaudhury

**Visualization:** Salma Haniffa, Michelle Ann Hughes, Aleksa Petković, Priyam Narain, Dipesh Chaudhury

**Funding acquisition:** Dipesh Chaudhury

**Investigation:** Vongai Mlambo, Salma Haniffa, Michelle Ann Hughes, Priyam Narain, Aleksa Petković, Marko Šušić

**Methodology:**Vongai Mlambo, Salma Haniffa, Michelle Ann Hughes, Priyam Narain

**Project administration:** Priyam Narain, Dipesh Chaudhury

**Resources:** Dipesh Chaudhury

**Software:** Salma Hanniffa, Michelle Ann Hughes, Aleksa Petković

**Validation:** Vongai Mlambo, Salma Haniffa, Michelle Ann Hughes, Priyam Narain, Aleksa Petković

**Writing-original draft:** Priyam Narain, Dipesh Chaudhury

**Writing-review-editing:** Aleksa Petković, Priyam Narain, Dipesh Chaudhury

**Supervision:** Dipesh Chaudhury

## Supplementary figures

**Supplemental Fig 1.**
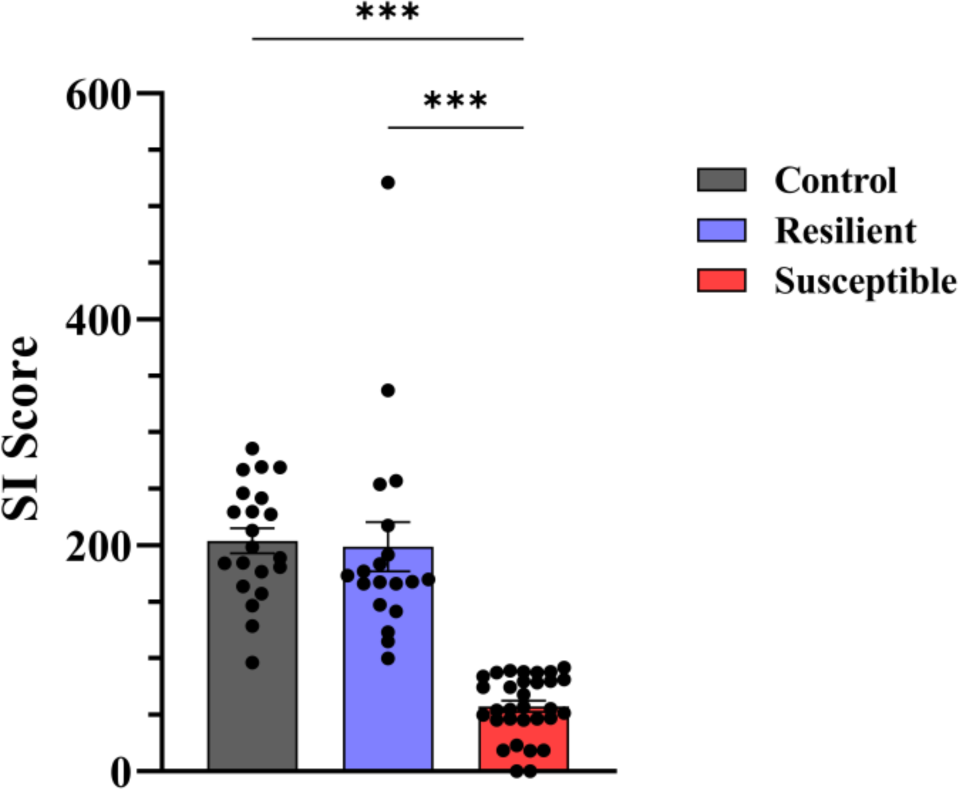
Aggregated social interaction (SI) scores following CSDS for all the control, resilient and susceptible mice utilized in the study. Experimental mice were exposed to the CSDS paradigm for 15 consecutive days. Following the final day of defeat, all mice were subjected to the SI test and the interaction scores were used to phenotype the mice as either stress susceptible or stress resilient. A one-way ANOVA revealed a significant difference in social interaction between control/ resilient and susceptible mice (F_2,68_ = 51.81, *p* = .07). There was a significant difference in SI values between control and susceptible (Tukey’s HSD, *p* < .001, 95% C.I. = [106.4, 186.5]) as well as resilient and susceptible mice (*p* < .001, 95% C.I. = [99.94, 182.5]). A total of *N* = 31 mice with SI <100 were identified to be stress susceptible. Control mice (*N* = 21) and stress resilient mice (*N* = 19) with SI >100 were selected for post-defeat molecular and recovery analysis. Tukey’s HSD test was used to compare the differences across phenotypes and error bars represent mean ± SEM. *** indicate *p* < .001.

**Supplemental Fig 2.**
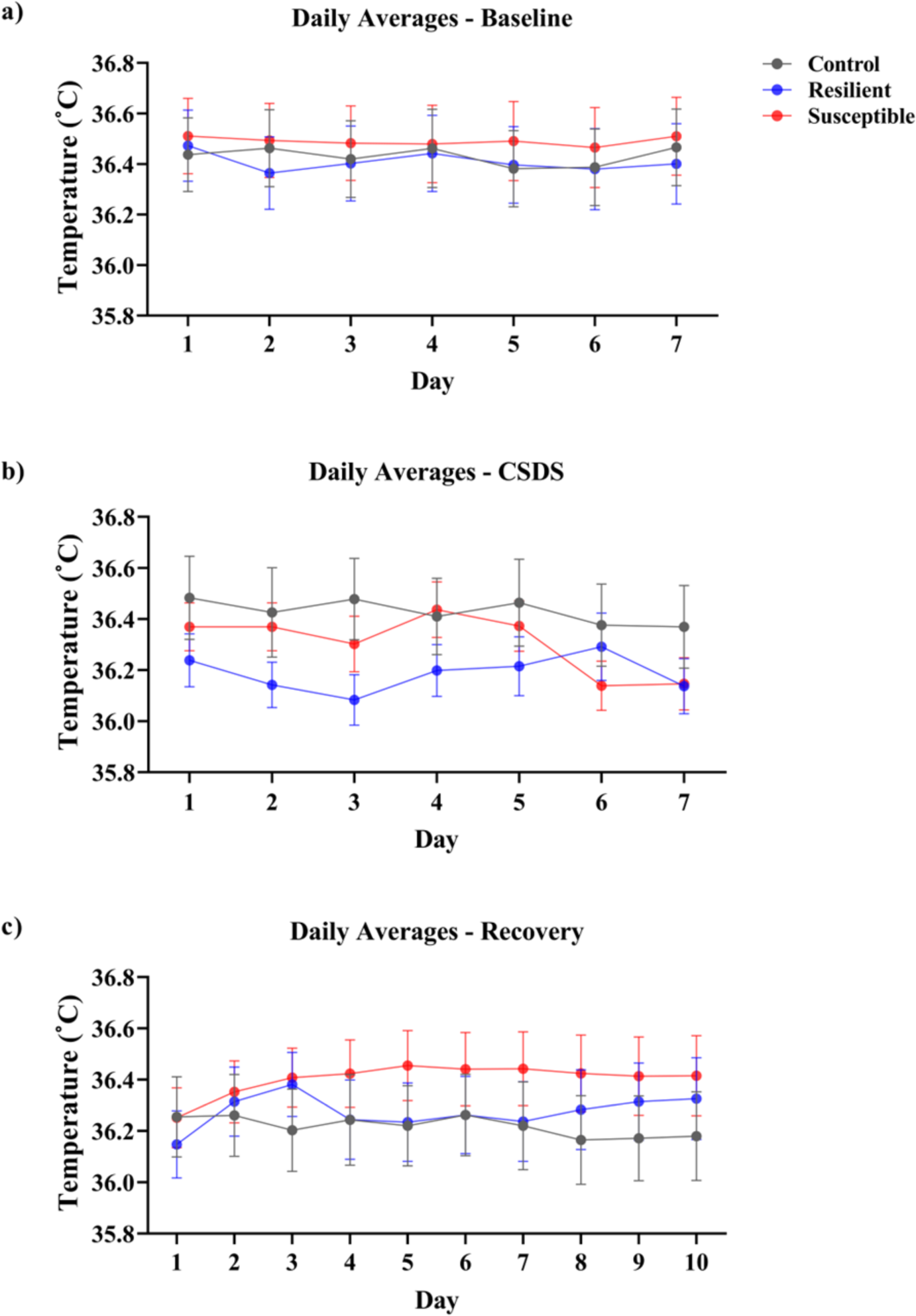
Daily temperatures averages: **(a)** The daily average temperature for each day was calculated for 7 days of baseline, **(b)** last 7 days of CSDS and **(c)** 10 days of post-CSDS recovery. Temperature was collected every hour. The error bars represent Mean ± SEM.

**Supplemental Fig 3.**
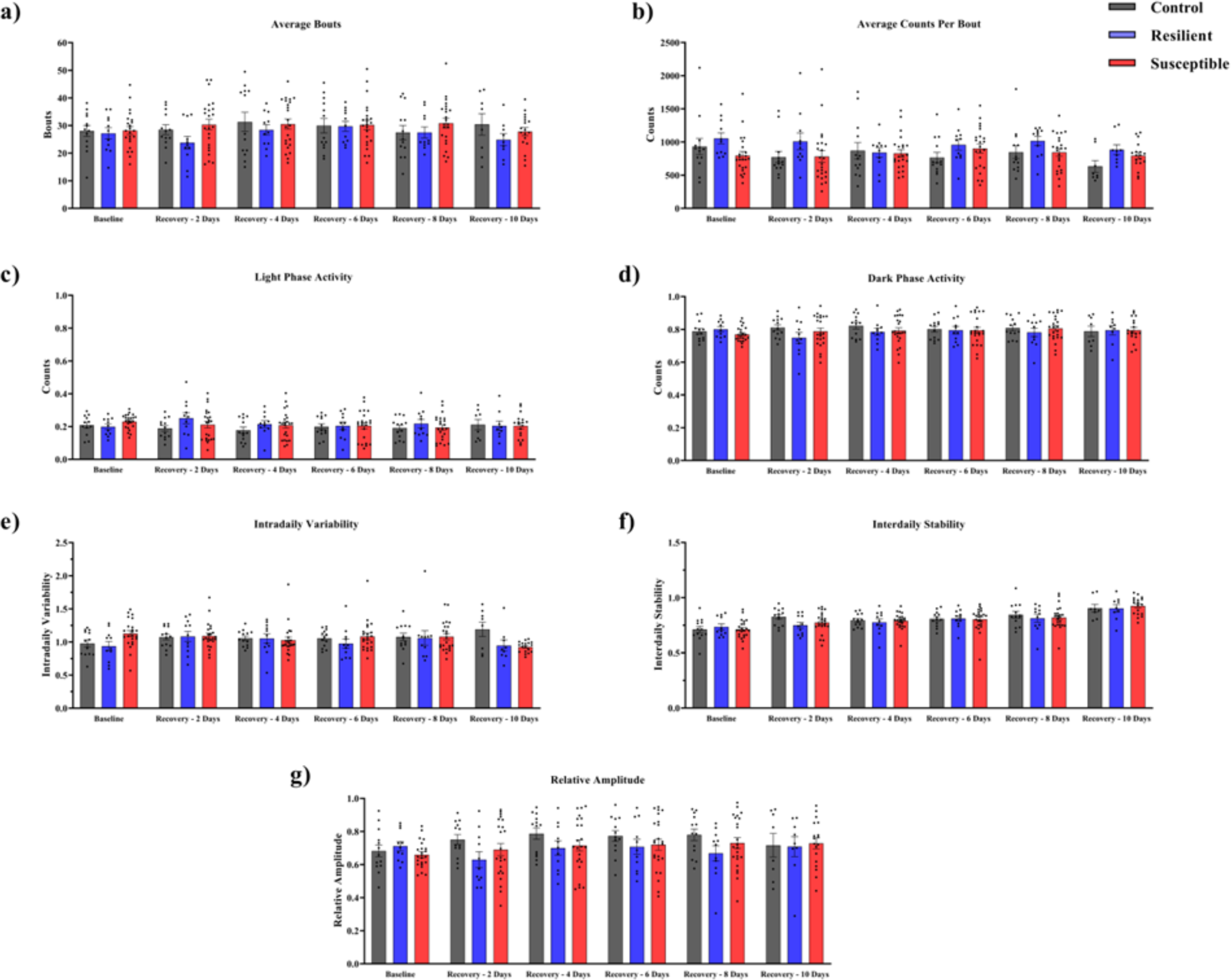
Chronic social stress has no effect on locomotor activity rhythms on resilient and susceptible mice following CSDS. **(a)** No significant difference in activity bouts. **(b)** No significant difference in activity counts. **(c & d)** Light phase and dark phase activity shows no significant difference between phenotypes. **(e & f)** No significant differences were observed in intrardaily variability and interdaily stability. **(g)** Relative activity amplitude showed no significant difference when comparing across phenotypes and conditions. **(a-g)** Control (*N* = 8 - 13), resilient (*N* = 9 - 11), and susceptible mice (*N* = 18 - 23). (e) Baseline is only shown for mice that underwent recovery. The error bars represent Mean ± SEM.

**Supplemental Fig 4.**
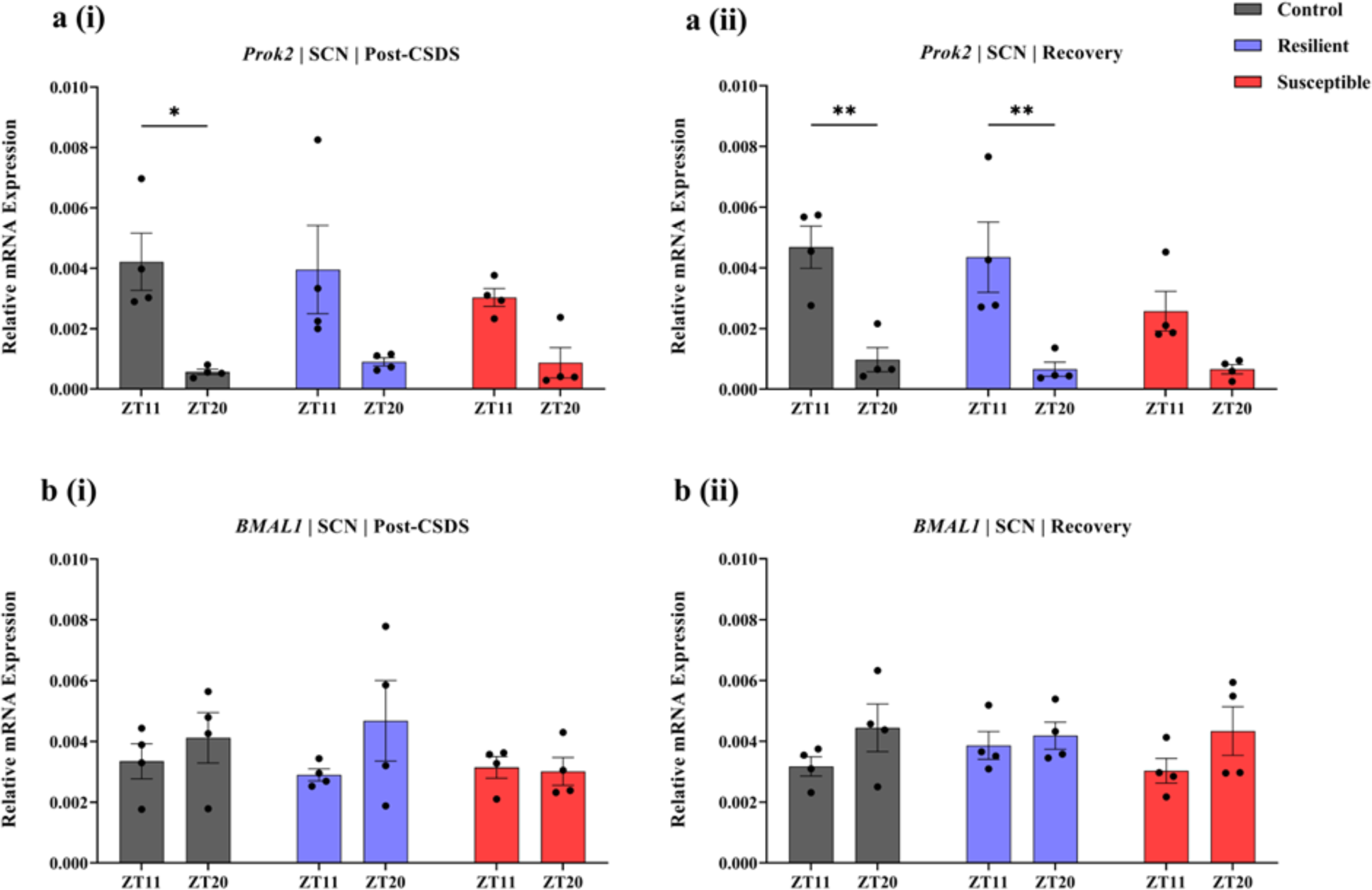
Expression analysis of clock genes *Prok2* and *BMAL1* in the SCN: **a(i, ii)**. Prok2 mRNA expression is similar in control, resilient and susceptible mice h post-CSDS and following recovery. **b(i, ii)**. There is no significant difference between the phenotypes in expression of clock gene *BMAL1*. Each experimental group consisted of 4 biological replicates split into 3 technical replicates (*N* = 4). Error bars represent mean ± SEM. * indicate *p* < .05, ** indicate *p* < .01.

**Supplemental Fig 5.**
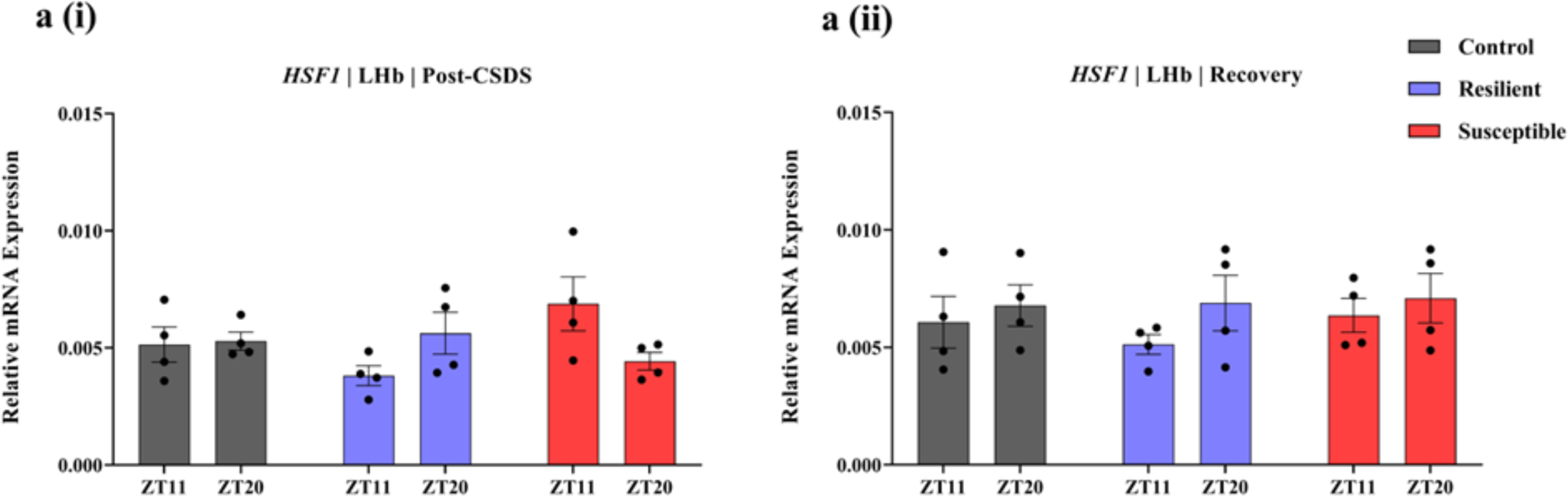
Expression analysis of *HSF1* in the LHb: **a(i,ii)** *Hsf1* mRNA levels does not show robust rhythmic expression in the LHb after CSDS or post recovery. Each experimental group consisted of 4 biological replicates split into 3 technical replicates (*N* = 4). Error bars represent mean ± SEM.

**Supplemental Fig 6.**
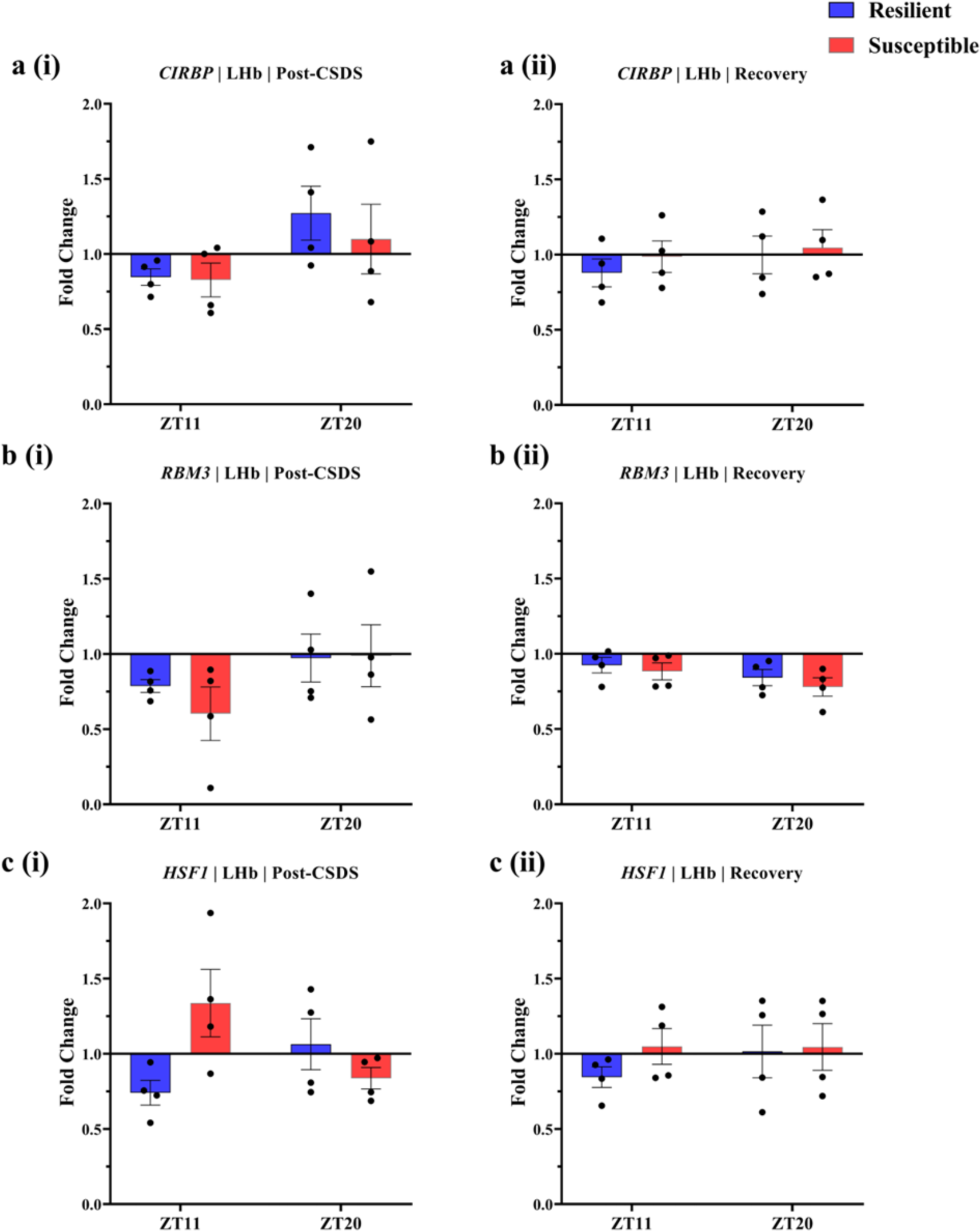
Normalized gene expression in the LHb of stress-exposed mice: Diurnal rhythmic expression of *Cirbp, RBM3* and *HSF1* in the LHb of resilient and susceptible mice normalized to controls immediately after CSDS and after 10 days of recovery. Each experimental group consisted of 4 biological replicates split into 3 technical replicates (*N* = 4).

**Supplemental Fig 7.**
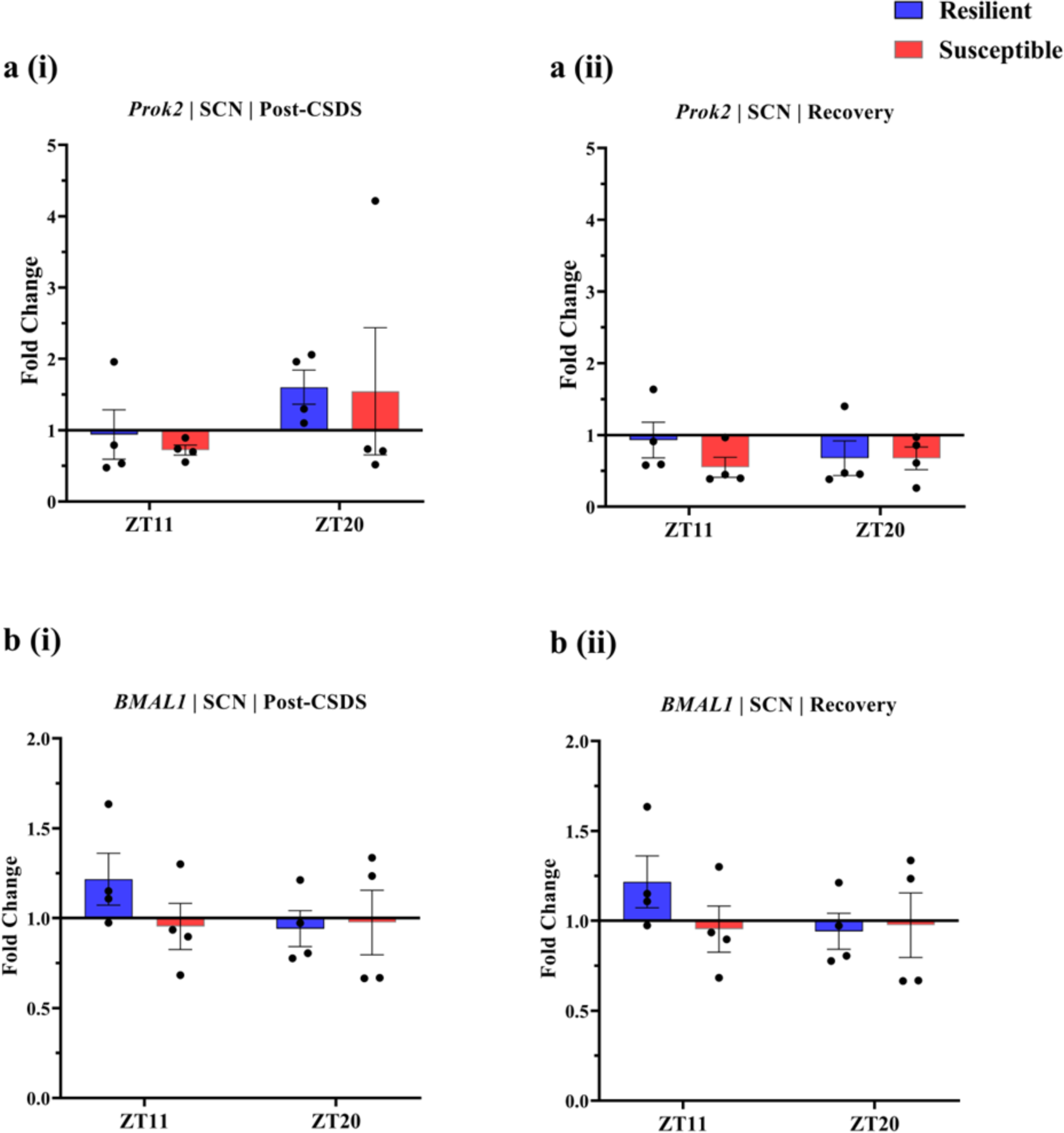
Normalized gene expression in the SCN of stress-exposed mice: Diurnal rhythmic expression of *Prok2* and *BMAL1* in the LHb of resilient and susceptible mice normalized to controls immediately after CSDS and after 10 days of recovery. Each experimental group consisted of 4 biological replicates split into 3 technical replicates (*N* = 4).

**Supplemental Fig 8.**
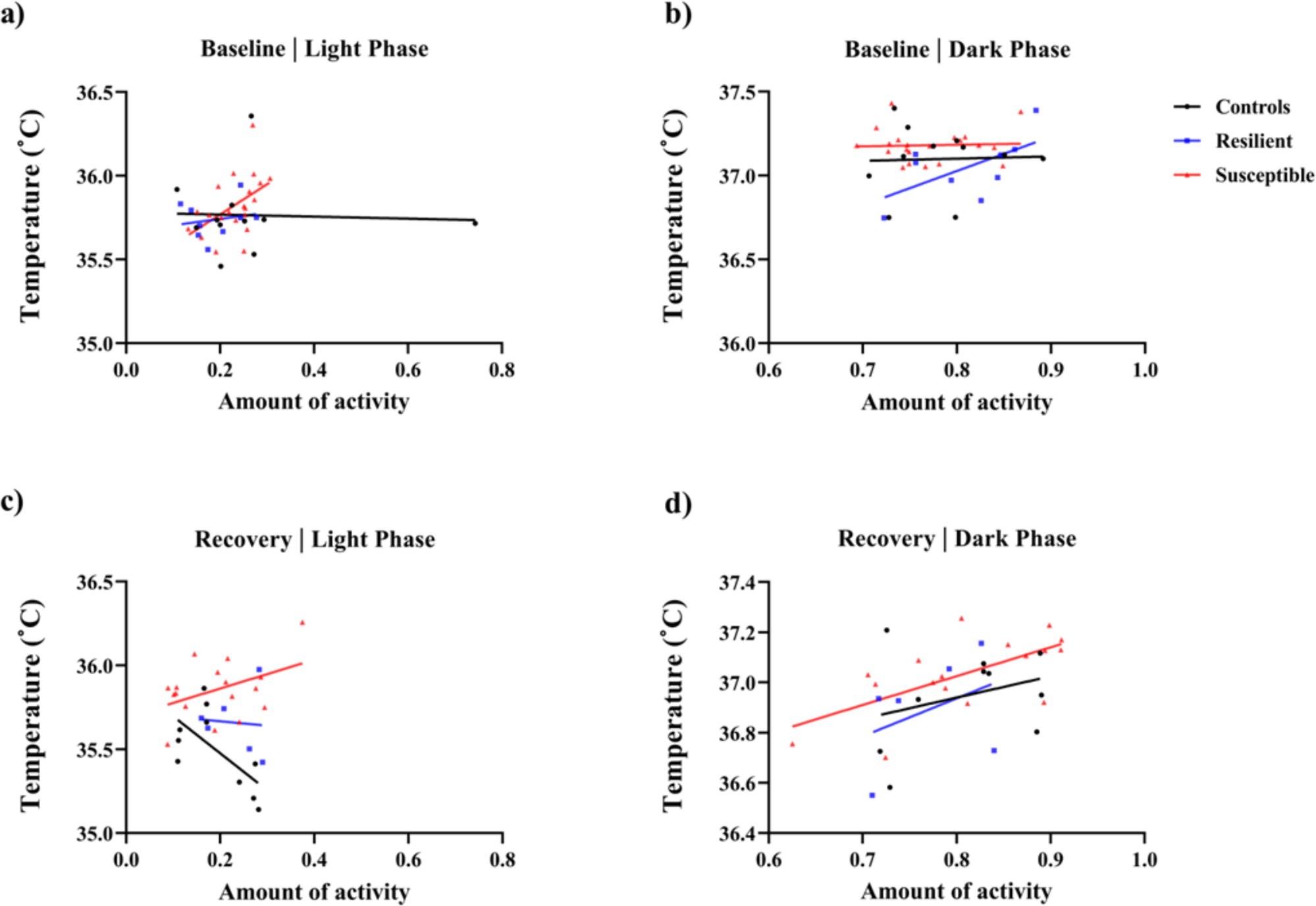
Linear correlation between temperature and locomotion activity counts: Average temperature across baseline/recovery was correlated with average amount of activity for each mouse. **(a)** During light phase at baseline, average temperature was significantly correlated with locomotion only in susceptible mice (r_1,19_ = 0.49, *p* = .03). **(b)** Temperature and dark-phase locomotor activity were not correlated in any of the phenotypes at baseline. **(c)** During recovery, stress-naive mice exhibited significant negative correlation between the temperature and amount of activity in light phase (r_1,8_ = 0.65, *p* = .04). **(d)** During dark phase in recovery, average temperature and locomotor activity were significantly correlated in susceptible mice (r_1,15_ = 0.64, *p* < .01). (a,b) Baseline: control (*N* = 11), resilient (*N* = 9), susceptible (*N* = 21). (c,d) Recovery: control (*N* = 10), resilient (*N* = 6), susceptible (*N* = 17).

**Table 1.**
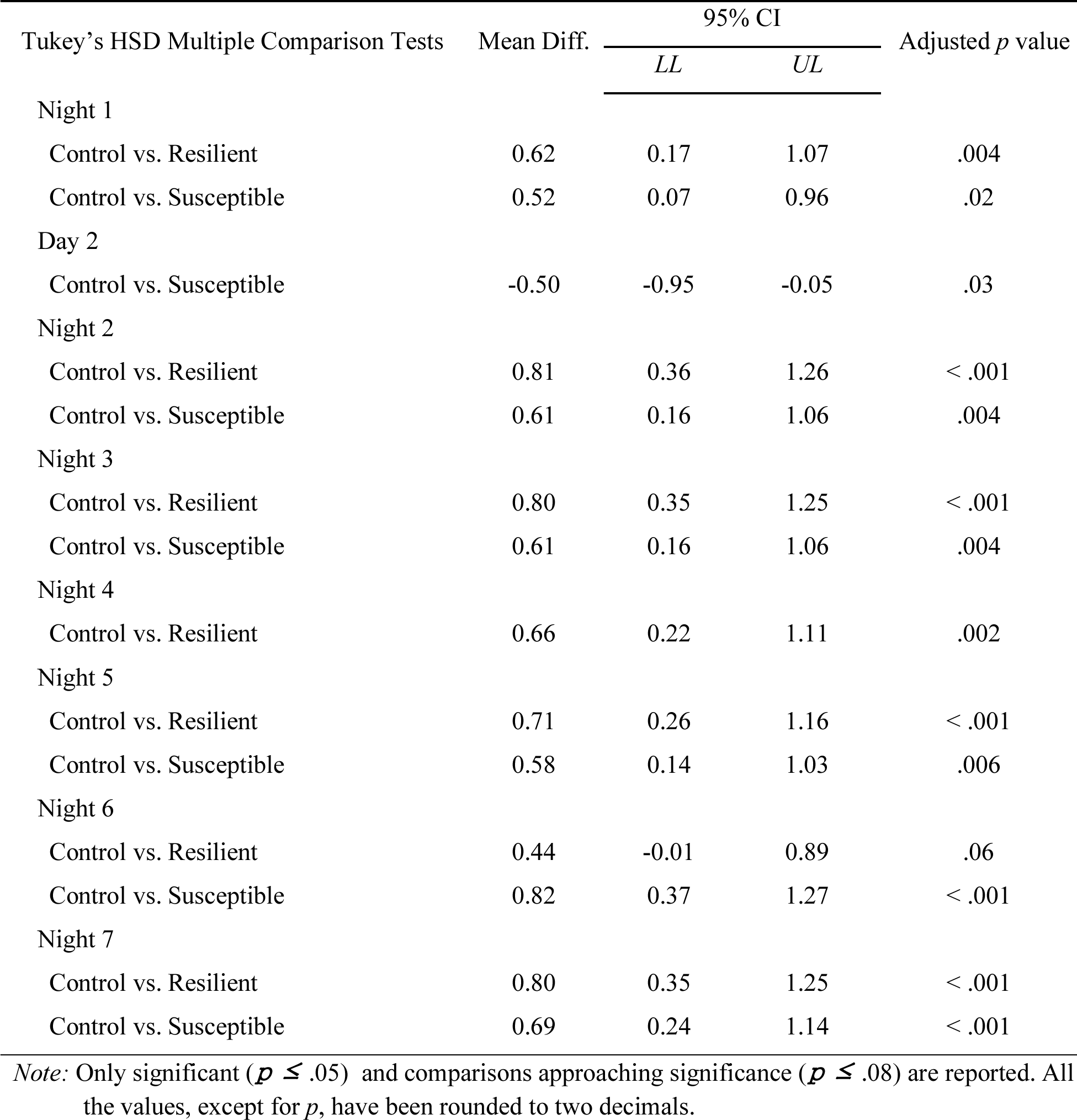
Tukey’s HSD multiple comparison tests for day and night average Tcore values during CSDS.

| Tukey's HSD Multiple Comparison Tests | Mean Diff. | 95% CI |  | Adjusted *p* value |
| --- | --- | --- | --- | --- |
|  |  | *LL* | *UL* |  |
| Night 1 |  |  |  |  |
| Control vs. Resilient | 0.62 | 0.17 | 1.07 | .004 |
| Control vs. Susceptible | 0.52 | 0.07 | 0.96 | .02 |
| Day 2 |  |  |  |  |
| Control vs. Susceptible | -0.50 | -0.95 | -0.05 | .03 |
| Night 2 |  |  |  |  |
| Control vs. Resilient | 0.81 | 0.36 | 1.26 | < .001 |
| Control vs. Susceptible | 0.61 | 0.16 | 1.06 | .004 |
| Night 3 |  |  |  |  |
| Control vs. Resilient | 0.80 | 0.35 | 1.25 | < .001 |
| Control vs. Susceptible | 0.61 | 0.16 | 1.06 | .004 |
| Night 4 |  |  |  |  |
| Control vs. Resilient | 0.66 | 0.22 | 1.11 | .002 |
| Night 5 |  |  |  |  |
| Control vs. Resilient | 0.71 | 0.26 | 1.16 | < .001 |
| Control vs. Susceptible | 0.58 | 0.14 | 1.03 | .006 |
| Night 6 |  |  |  |  |
| Control vs. Resilient | 0.44 | -0.01 | 0.89 | .06 |
| Control vs. Susceptible | 0.82 | 0.37 | 1.27 | < .001 |
| Night 7 |  |  |  |  |
| Control vs. Resilient | 0.80 | 0.35 | 1.25 | < .001 |
| Control vs. Susceptible | 0.69 | 0.24 | 1.14 | < .001 |

*Note:* Only significant ( $p \leq .05$ ) and comparisons approaching significance ( $p \leq .08$ ) are reported. All the values, except for $p$ , have been rounded to two decimals.

**Table 2.** Tukey’s HSD multiple comparison tests for Tcore amplitude values.

| Tukey's HSD Multiple Comparison Tests | Mean Diff. | 95% CI |  | Adjusted *p* value |
| --- | --- | --- | --- | --- |
|  |  | *LL* | *UL* |  |
| CSD |  |  |  |  |
| Control vs. Resilient | 0.5 | 0.29 | 0.71 | < .001 |
| Control vs. Susceptible | 0.45 | 0.27 | 0.63 | < .001 |
| Recovery (Day 1 – Day 5) |  |  |  |  |
| Control vs. Resilient | 0.19 | -0.03 | 0.41 | .17 |
| Control vs. Susceptible | 0.28 | 0.09 | 0.47 | < .001 |
| Recovery (Day 6 – Day 10) |  |  |  |  |
| Control vs. Resilient | 0.14 | -0.08 | 0.36 | .61 |
| Control vs. Susceptible | 0.15 | -0.04 | 0.34 | .26 |

*Note:* All the values, except for *p*, have been rounded to two decimals.

